# SARS-COV-2 nucleocapsid protein hijacks multiple components of the host nuclear transport machinery for distinct functions

**DOI:** 10.1101/2024.08.09.607288

**Authors:** Angela R. Harrison, Sui Z. Wang, Kylie M. Wagstaff

## Abstract

Many viruses target the host nuclear transport machinery to traffic their own proteins or restrict trafficking of host cargo, including mediators of antiviral immune signalling. Nucleocapsid (N) proteins from several coronaviruses traffic to the nucleus/nucleolus, with roles in cell cycle regulation. While N-protein from severe acute respiratory syndrome virus 2 (SARS-COV-2), the causative agent of the global COVID-19 pandemic, is widely reported to localise to the cytoplasm, we identify multiple, functionally distinct interactions of N-protein with the host’s nuclear transport machinery. Using quantitative cell imaging, including fluorescence recovery after photobleaching, and protein-protein interaction analysis, we describe a sub-population of SARS-COV-2 N-protein that localises more diffusely between the nucleus and cytoplasm, undergoes active nuclear import, and re-localises nuclear import receptors (karyopherins) KPNB1 and KPNA2 to the nucleus and cytoplasm, respectively. Truncation analyses identify at least two distinct KPNA2/KPNB1 binding sites located in the N-terminal and C-terminal regions of N-protein. Interestingly, while mutation of K/R-rich sites within these domains reduces KPNA2/KPNB1 binding and disables re-localisation of KPNA2, re-localisation of KPNB1 and nuclear import of N-protein remain intact, indicating that these are molecularly and functionally distinct mechanisms. siRNA knockdown confirms a role for KPNB1 in N-protein nuclear trafficking, while KPNA2 binding and mislocalisation may be antagonistic. Thus, SARS-COV-2 N-protein binds to karyopherins *via* multiple distinct sites to facilitate import and other functions.

## Introduction

Severe acute respiratory syndrome coronavirus 2 (SARS-COV-2) is responsible for the COVID-19 global pandemic, causing more than 775 million infections and seven million deaths worldwide [1]. While we now have access to vaccines and a limited number of antivirals, the continuing emergence of new SARS-COV-2 variants reduce their effectivity, highlighting the need for ongoing research to develop additional therapeutic strategies.

Like other coronaviruses, SARS-COV-2 has a positive-sense RNA genome that is replicated on modified membranes within the cytoplasm. Viruses that are replicated exclusively in the cytoplasm, including coronaviruses, often traffic proteins to the host cell nucleus to mediate ‘accessory’ functions, and these represent an attractive target for antivirals [2]. All molecules entering or exiting the nucleus must pass through nuclear pore complexes (NPCs) embedded in the nuclear envelope (reviewed in [3]). Molecules < c. 40 kDa can passively diffuse through NPCs; however, this is inefficient, and so many proteins, including proteins larger than the diffusion limit, engage nuclear transport receptors of the karyopherin (KPN) superfamily for active trafficking, a highly regulated, energy-dependent process. KPNs recognise nuclear localisation and nuclear export sequences (NLS and NES, respectively) in cargo proteins. In the classical import pathway, an NLS consisting of a single or bi-partite stretch of positively-charged residues is recognised in the cytoplasm by a KPN alpha (KPNA) ‘adapter’. KPNA also engages KPN beta 1 (KPNB1), which facilitates movement of the trimeric complex through the NPC. In the nucleus, binding of Ran-GTP to KPNA/KPNB1 releases the cargo into the nucleoplasm. KPNB1 can also mediate nuclear import independently of KPNA *via* binding to certain cargo directly. To return to the cytoplasm, nuclear cargoes typically engage Ran-GTP-bound “exportin” KPNs *via* distinct NES sequences; cargoes are trafficked through NPCs and then released into the cytoplasm upon GTP hydrolysis by cytosolic enzymes [3].

Diverse viruses hijack the nuclear transport machinery to transport their own proteins [2,3]. Nucleocapsid (N) proteins from many coronaviruses encode NLS and NES sequences to shuttle between the cytoplasm, where they encapsidate viral RNA during virus replication and assembly, and the nucleus/nucleolus (reviewed in [4]). For infectious bronchitis coronavirus (IBV), nuclear/nucleolar N-protein regulates cell cycle progression [5]. While N-protein from SARS-COV-2 appears predominantly cytoplasmic [6–10], it shares c. 90% sequence identity with SARS-COV-1 [11], which has several reported NLSs [4,12–14]. In fact, recent studies have reported localisation of SARS-COV-2 N-protein to the nucleus and/or nucleoli in infected [15] and transfected [16–18] cells, indicating that nuclear trafficking is likely conserved. However, this has not been directly examined. Many viruses also target these pathways to inhibit trafficking of host cargo, which can in turn mediate suppression of antiviral immune responses [3]. For example, ORF6 from SARS-COV-1 binds KPNA2 and KPNB1, tethering the complex to the ER to inhibit nuclear import of STAT1 [19], a key mediator of antiviral interferon signalling. A similar mechanism appears to be used by SARS-COV-2 [20].

Here, we investigated nuclear trafficking of SARS-COV-2 N-protein, identifying a sub-population that undergoes nuclear import. We also identify two separate sites within N-protein that bind to KPNA2/KPNB1; however, these do not appear to contribute to N-protein nuclear import, but rather mediate mislocalisation of KPNAs. The data suggest that a distinct, KPNB1-dependent mechanism/pathway drives N-protein nuclear trafficking.

## Results

### SARS-COV-2 N-protein is imported into the nucleus in a proportion of cells

N-proteins from many coronaviruses, including SARS-COV-1 and IBV, traffic to the host cell nucleus/nucleolus [4,12–14]. While SARS-COV-2 N-protein is largely reported to localise to the cytoplasm [6–10], it has been observed within the nucleus and/or nucleolus in transfected and infected cells [15–18], indicative of nuclear trafficking. We therefore assessed the localisation of SARS-COV-2 N-protein over time using confocal laser scanning microscopy (CLSM). HCT-8 cells, which are permissible to SARS-COV-2 infection, were transfected to express GFP-tagged N-protein (GFP-N) for 6 to 72 h before fixation and imaging (Figure 1A). Nucleocytoplasmic localisation was then quantified by calculating the nuclear to cytoplasmic fluorescence ratio (Fn/c; Figure 1B), as previously [21–23]. GFP-N was predominantly excluded from the nucleus at all time points (Fn/c < 0.5). Interestingly, GFP-N localised more diffusely between the nucleus and cytoplasm in a proportion of cells (Fn/c > 0.5); this sub-population (termed ‘diffuse’) was most apparent at 16 h and 24 h post-transfection, representing 30-45% of cells (Figure 1C), and least apparent at 72 h, representing only 3% of cells, and largely accounted for the variability in mean Fn/c values between timepoints (Figure 1B). Since nucleolar localisation of IBV N-protein is affected by the cell cycle [24], we considered that the distinct populations of SARS-COV-2 N-protein may represent cells in different cell states; however, the distinct populations were still present after cell synchronisation with a similar trend over time (Figure S1). Thus, N-protein localisation appears to be variable and dynamic, with N-protein localising in the nucleus under unknown cellular conditions. Importantly, N-protein appeared excluded from nucleoli at all time points examined (Figure 1 and Figure S1), in contrast to N-protein from other coronaviruses [4], indicative of virus-specific localisation mechanisms.

**Figure 1.**
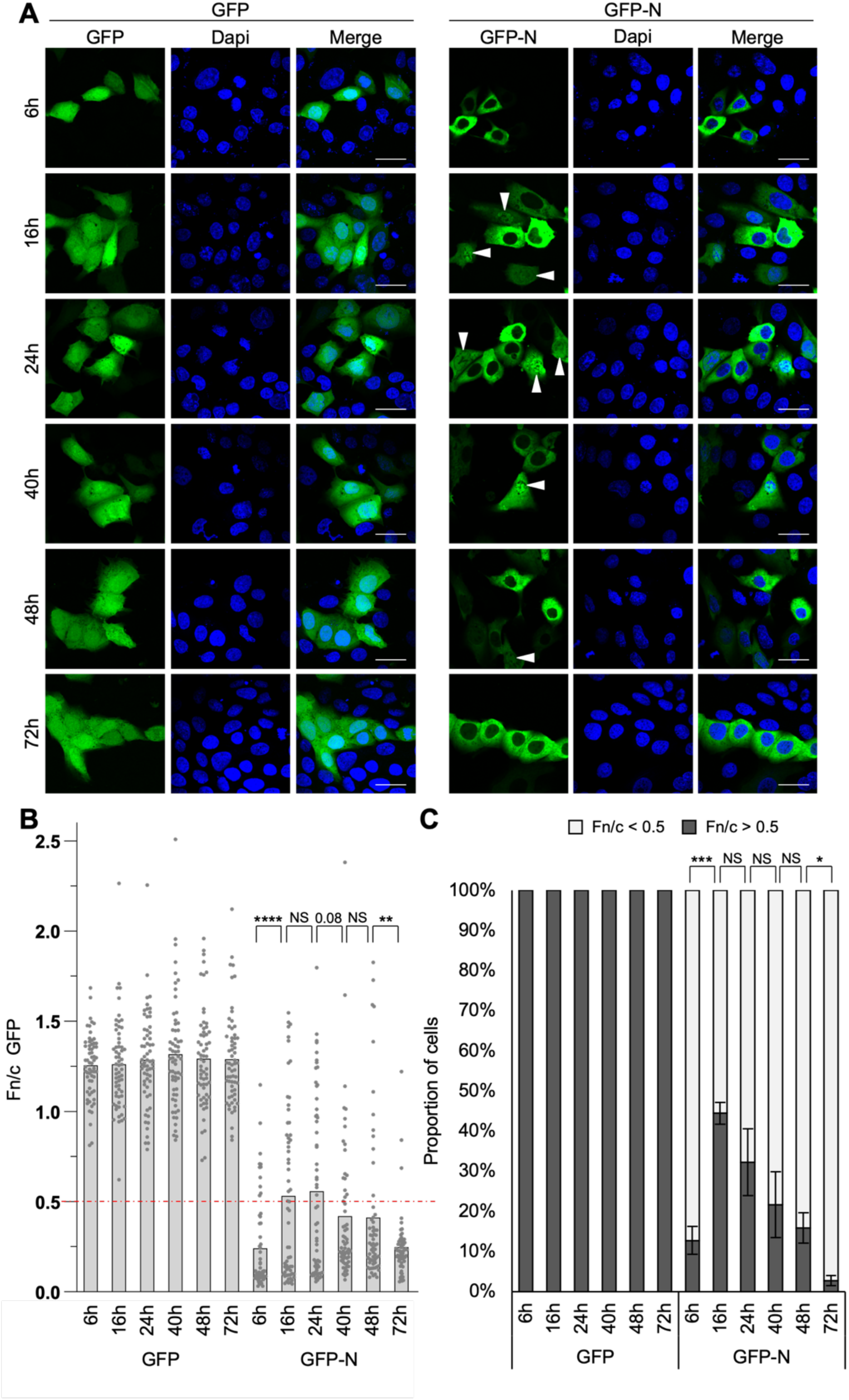
Localisation of SARS-COV-2 N-protein changes over time. (A) HCT-8 cells were transfected to express GFP or GFP-N for the indicated time before fixation and CLSM analysis. Representative images are shown. (B,C) Images such as those in (A) were analysed to calculate the Fn/c for GFP (B; mean, n = 62 cells for each condition from a representative assay) and the percentage of cells with Fn/c < or > 0.5 (C; mean ± SEM, n = 3 independent assays). Dapi (blue) was used to localise nuclei. Arrows indicate cells with Fn/c for GFP-N > 0.5. Statistical analysis used Student’s *t*-test. *, p < 0.05; **, p < 0.01; ***, p < 0.001; ****, p < 0.0001; NS, not significant. Scale bars, 30 μm.

Given that N-protein with a GFP tag is > 70 kDa, exceeding the NPC diffusion limit, nuclear localisation is indicative of an active nuclear import mechanism being present. To examine the kinetics of SARS-COV-2 N-protein nuclear import, we next performed fluorescence recovery after photobleaching (FRAP) on transfected HCT-8 cells. Briefly, GFP-N in the nucleus was bleached using high-powered laser and the nuclear fluorescence recovery was monitored over time by CLSM (Figure 2A). GFP, which is small enough to diffuse between the nucleus and cytoplasm, was used as a positive control for nuclear entry, and GFP nuclear fluorescence consistently and rapidly recovered after bleaching, as expected (Figure 2B). In contrast, the responses in GFP-N-expressing cells were varied, with GFP-N able to recover in some cells, consistent with nuclear import, but not others. Importantly, the capacity of GFP-N to recover correlated with its pre-bleach localisation (Figure 2C); cells with a pre-bleach Fn/c > 0.5 (‘diffuse’) had a dramatically higher initial rate of recovery (Figure 2D) and maximal recovery (Figure 2E) than cells with a pre-bleach Fn/c < 0.5 (‘cytoplasmic’). This suggests that the distinct populations of N-protein reflect differences in nuclear import activity, with the diffuse sub-population mobile and capable of nuclear entry and the cytoplasmic sub-population less so.

**Figure 2.**
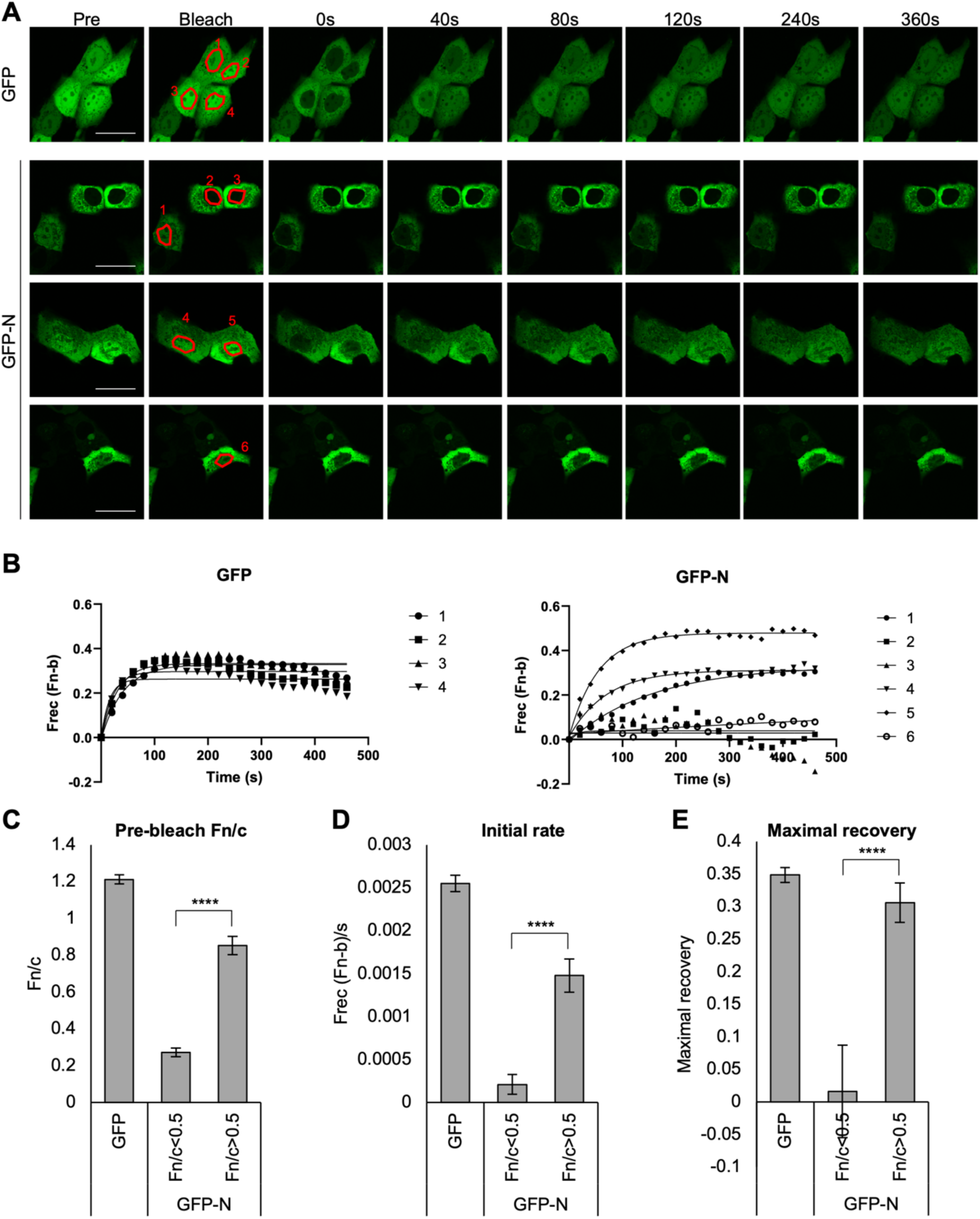
SARS-COV-2 N-protein undergoes nuclear import. (A) HCT-8 cells transfected to express GFP or GFP-N were imaged by CLSM prior (Pre) to photobleaching in the indicated nucleus (Bleach) and then monitored every 20s for 460s. Representative images are shown. (B) Digitised images, such as those in (A), were analysed to determine the fractional change of nuclear fluorescence (Frec(Fn-b)). (C) The pre-bleach Fn/c value was determined and GFP-N samples were grouped based on Fn/c < or > 0.5. (D, E) Curves such as those in B were used to determine the initial rate of import (initial 100s post-bleaching, D) and maximal recovery of nuclear fluorescence (E). Bar charts represent the mean ± SEM (n ζ 29 cells for each condition). Statistical analysis used Student’s *t*-test. ****, p < 0.0001. Scale bars, 30 μm.

### SARS-COV-2 N-protein associates with and re-localises the nuclear transport machinery

Many host and viral cargoes utilise the classical KPNA/KPNB1 pathway for nuclear import [2,3]. We thus examined whether SARS-COV-2 N-protein can associate with KPNs by co-immunoprecipitation from transfected HCT-8 cells (Figure 3A). GFP fused to the NLS from the simian virus 40 (SV40) large tumour antigen (T-ag) protein was used as a positive control, as previously [25,26]. GFP-T-ag_NLS_ co-precipitated endogenous KPNA2, as expected, although the interaction was difficult to detect, consistent with the transient nature of trafficking complexes. T-ag_NLS_ did not detectably co-precipitate KPNB1, again likely due to poor retention of the transient complex during lysis and immunoprecipitation; KPNA2 acts as the adaptor between KPNB1 and T-ag_NLS_ and is therefore more likely to be detected. Interestingly, GFP-N clearly co-precipitated both endogenous KPNA2 and KPNB1, well above that observed for T-ag_NLS_, indicative of a robust interaction (Figure 3A). Similar results were observed from cells over-expressing FLAG-tagged KPNA2 (Figure S2A).

**Figure 3.**
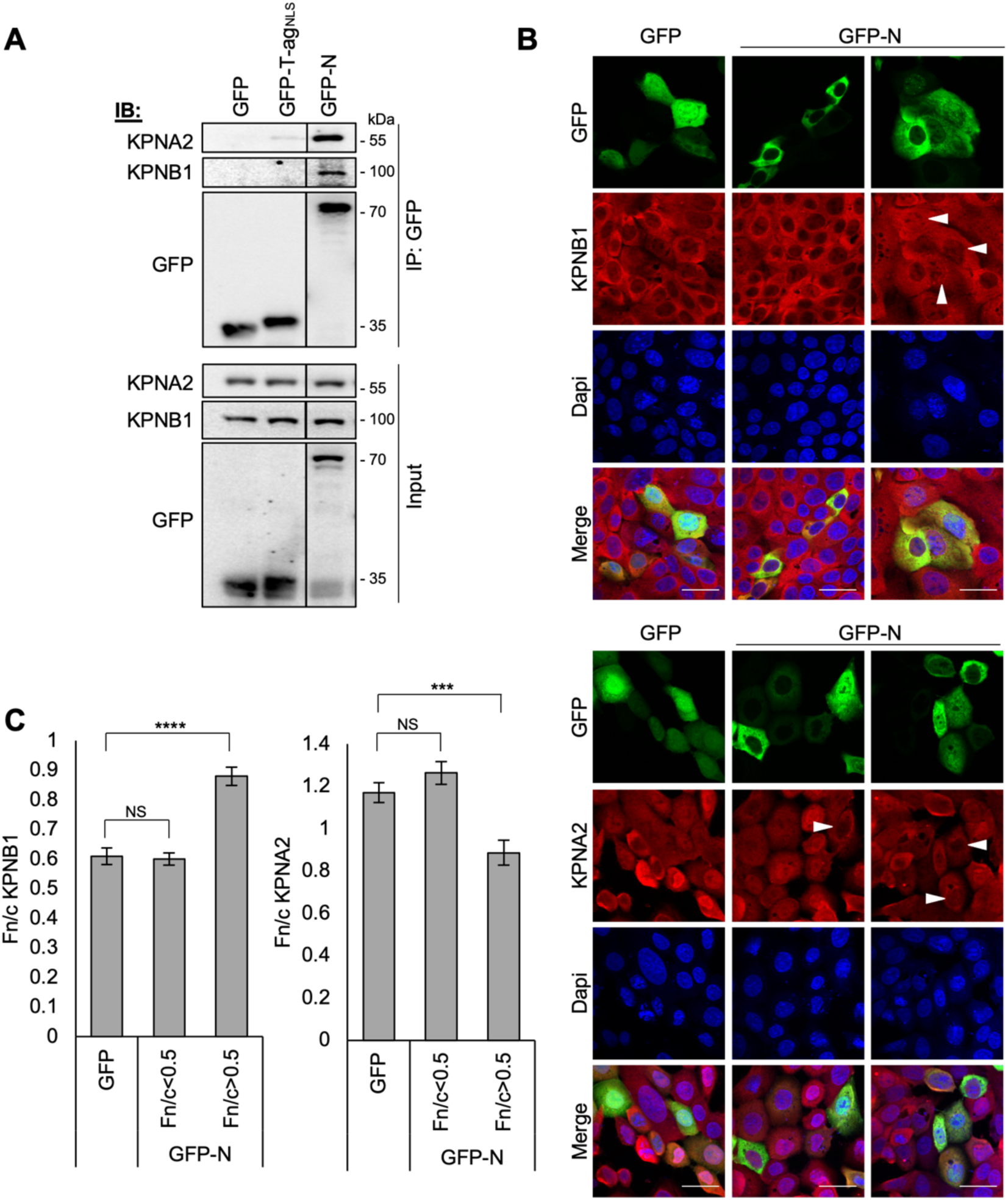
SARS-COV-2 N-protein associates with and re-localises KPNB1 and KPNA2. (A) HCT-8 cells were transfected to express GFP, GFP-T-ag_NLS_ or GFP-N for 24 h before lysis and immunoprecipitation for GFP. Lysates (Input) and immunoprecipitates (IP) were analysed by immunoblotting (IB) using antibodies against the indicated proteins. Data are from a single blot with intervening and marker lanes removed. (B,C) HCT-8 cells were transfected to express the indicated proteins for 16 h before fixation, immunofluorescent staining (red) for KPNB1 (upper panel) or KPNA2 (lower panel) and CLSM (B) to determine the Fn/c for KPNB1 and KPNA2 (C; mean ± SEM, n = 62 cells for each condition). Only cells with clear expression of GFP or GFP-N protein were analysed. GFP-N data were grouped based on GFP-N Fn/c values (Fn/c < or > 0.5). Dapi (blue) was used to localise nuclei. Arrows indicate cells with Fn/c for GFP-N > 0.5. Statistical analysis used Student’s *t*-test. ***, p < 0.001; ****, p < 0.0001; NS, not significant. Scale bars, 30 μm.

Although it appears likely that the interaction of SARS-COV-2 N-protein with KPNA2/KPNB1 facilitates nuclear import, viral proteins do not always associate with the trafficking machinery for their own transport. ORF6 from SARS-COV-1, for example, tethers KPNA2/KPNB1 complexes to the ER/golgi to inhibit nuclear import of the immune signalling mediator STAT1 [19]. We therefore assessed the effect of N-protein expression on KPNA2/KPNB1 localisation by CLSM (Figure 3B-C). KPNB1 was detected in both the cytoplasm and nucleus of GFP-expressing control cells, moderately favouring cytoplasmic localisation (Fn/c ∼0.6), but became significantly more nuclear (Fn/c ∼0.9) in the presence of ‘diffuse’ GFP-N, suggesting that the KPNB1-N association can drive nuclear import of the complex. Importantly, highly cytoplasmic GFP-N had no effect on KPNB1 localisation, consistent with the lack of nuclear entry of this population (Figure 2). This also indicates that, unlike SARS-COV-1-ORF6, SARS-COV-2 N-protein does not associate with KPNB1 to sequester the transporter in the cytoplasm.

Surprisingly, analogous analysis of KPNA2 localisation indicated a consistent but inverse effect; KPNA2 localised more diffusely between the nucleus and cytoplasm in control cells (Fn/c ∼1.2), and the diffuse population of GFP-N effected a significant decrease in KPNA2 nuclear localisation (Fn/c ∼0.9), while cytoplasmic GFP-N had no effect (Figure 3C). This is opposite to the effect of T-ag_NLS_, which traffics *via* KPNA2 and promotes KPNA2 nuclear accumulation when expressed (Figure S3), but similar to SARS-COV-2 ORF6, which re-localises KPNA2 to the cytoplasm (Figure S3) and inhibits nuclear transport [20,27,28]. Thus, N-protein associations with KPNB1 and KPNA2 may have distinct functions.

### SARS-COV-2 N-protein binds KPNB1 and KPNA2 via multiple domains

Given the apparently opposing effects of SARS-COV-2 N-protein on KPNA2 and KPNB1 localisation, we next sought to further define the associations. KPNA/KPNB1 heterodimers commonly recognise ‘classical’ NLS sequences, comprising one or more short strings of positively charged residues. Inspection of the N-protein amino acid sequence identified several NLS-like sequences throughout the protein, many of which are conserved among coronaviruses [29]. N-protein is a multi-domain protein, comprising structured N- and C-terminal domains connected by a S/R-rich flexible linker region (LKR), and bookended by short N- and C-terminal tails (Figure 4A). To locate the KPNA2/KPNB1 binding site/s, we generated various constructs to express truncated GFP-N proteins (Figure 4A). The truncations were assessed for their steady state localisation (Figure S4) and capacity to bind KPNA2 and KPNB1 (Figure 4B). GFP-N_1-174_, which includes an NLS-like sequence at residues 88-95, strongly co-precipitated KPNA2/KPNB1, indicative of a binding site in the N-arm/NTD region. Addition of the LKR (GFP-N_1-274_) had no discernible effect on binding. GFP-N_175-364_ also clearly co-precipitated KPNA2/KPNB1, suggesting that there is an additional binding site in the N-protein LKR/CTD. This appears to require both the LKR and CTD, as no or minimal KPNA2/KPNB1 was co-precipitated by the LKR (GFP-N_175-247_) or CTD (GFP-N_248-364_) on their own. There is an NLS-like sequence at residues 256-262 (Figure 5A), which is within the CTD but close to the LKR junction, such that both domains may be required for the overall structure and accessibility/presentation of this NLS. Notably, addition of the C-tail to the LKR-CTD fragment (GFP-N_175-419_) enhanced KPNA2/KPNB1 binding (Figure 4B), identifying a possible auxiliary role for the C-tail. However, it seems unlikely that this region is the main driver of KPNA2/KPNB1 interaction, given the lack of binding by the C-tail alone (GFP-N_365-419_), and minimal binding by the CTD-C-tail fragment (GFP-N_248-419_).

**Figure 4.**
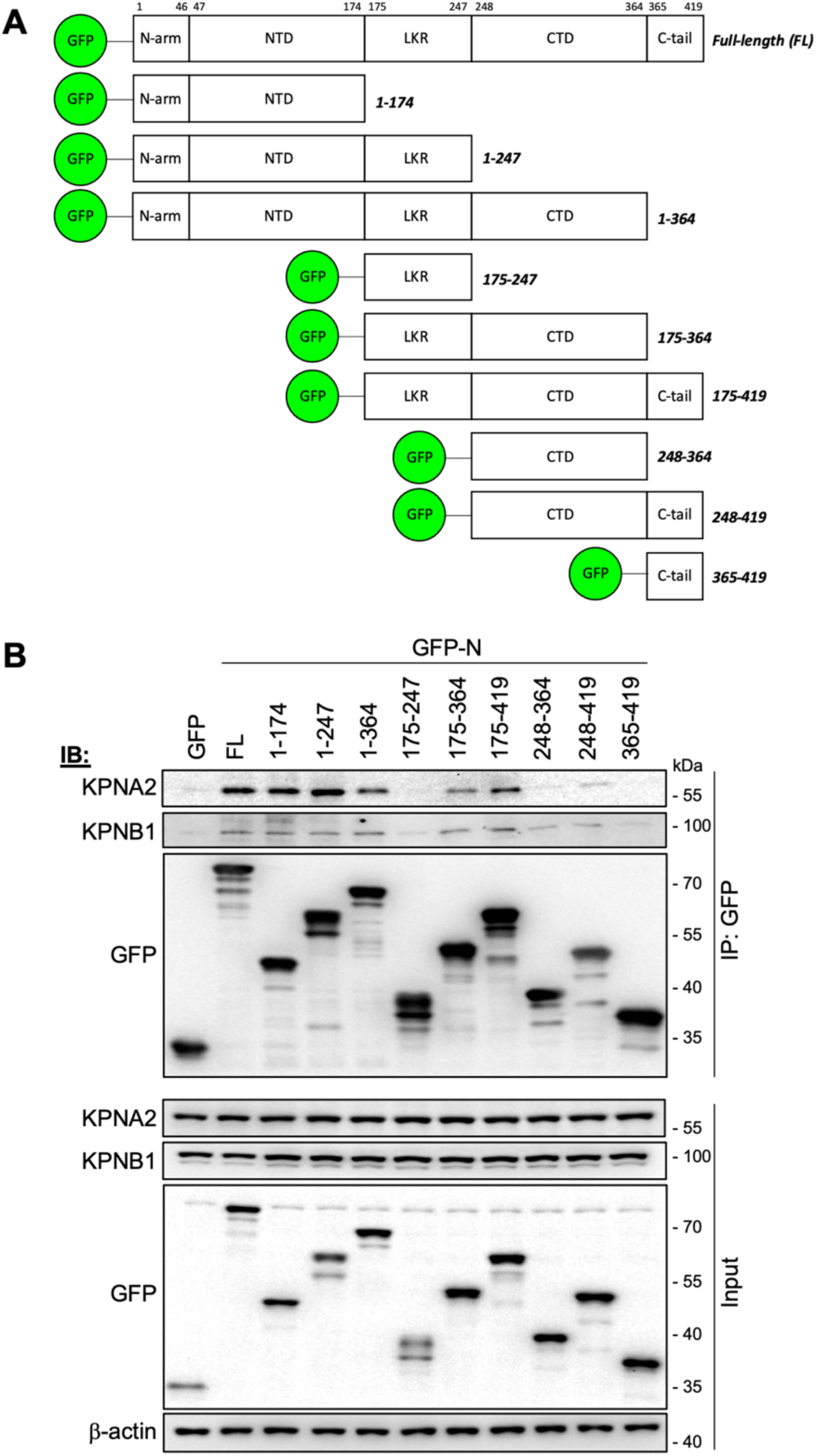
SARS-COV-2 N-protein interacts with KPNs through multiple binding regions. (A) Schematic of SARS-COV-2 N-protein truncations generated in this study. Full-length (FL) N-protein consists of structured N-terminal (NTD) and C-terminal (CTD) domains interspersed between an N-arm, C-tail and central linker region (LKR). Numbering indicates residue positions in FL N-protein. (B) HCT-8 cells were transfected to express the indicated proteins for 16 h before lysis and immunoprecipitation for GFP. Lysates (Input) and immunoprecipitates (IP) were analysed by immunoblotting (IB) using antibodies against the indicated proteins. Results are representative of two independent assays.

**Figure 5.**
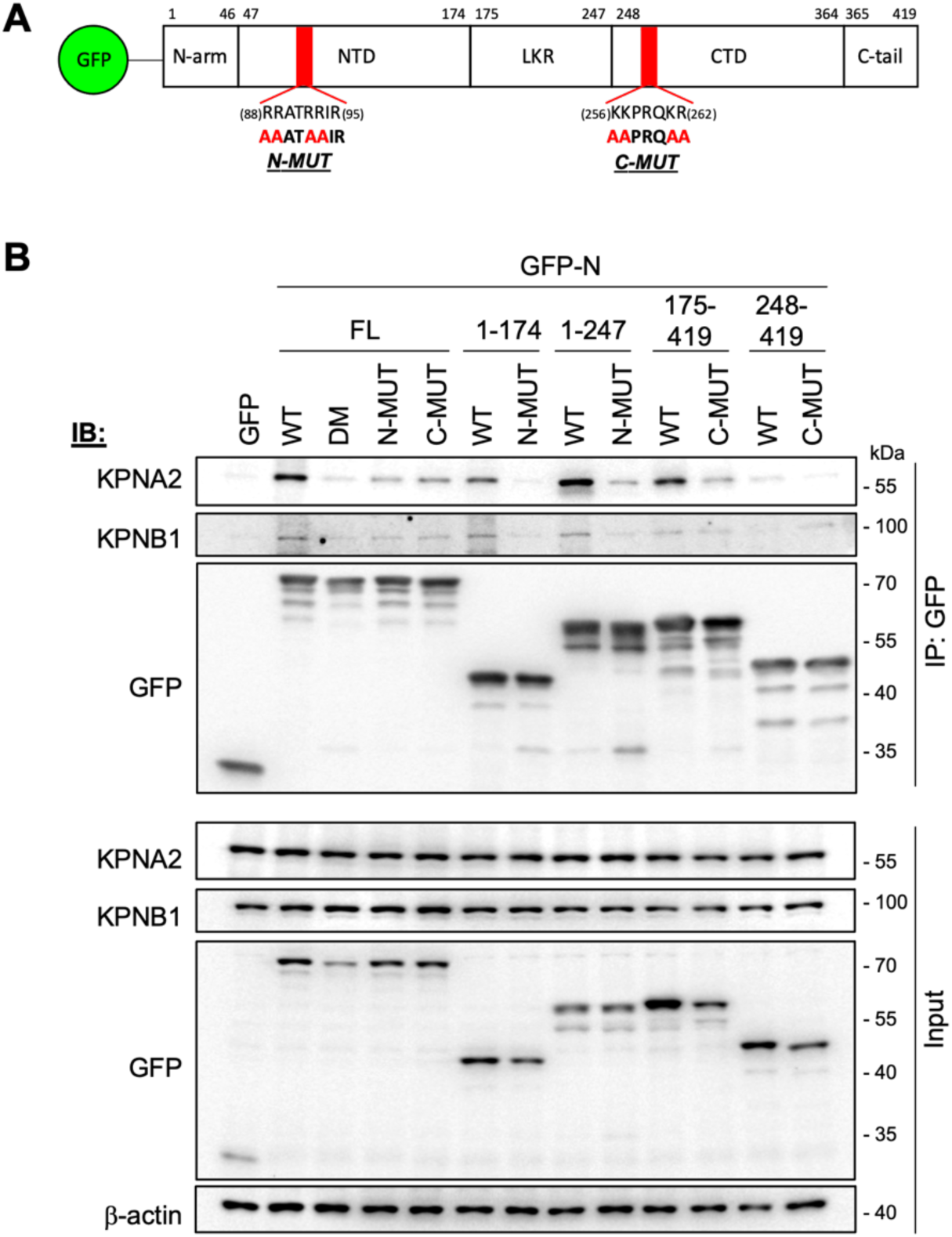
Mutation to NLS-like sequences in the CTD and NTD of SARS-COV-2 N-protein impairs binding to KPNs. (A) Schematic of N-protein domains as described in the legend to Figure 4A. Mutations to NLS-like sequences in the NTD (N-MUT) and CTD (C-MUT) are indicated. (B) HCT-8 cells were transfected to express the indicated proteins for 16 h before lysis, immunoprecipitation for GFP, and IB, as described in the legend to Figure 4B. Double mutant (DM) contains N-MUT and C-MUT. Results are representative of two independent assays.

Based on the truncation results, we next disabled potential NLS-like sequences in the NTD and CTD by mutating key R/K residues to alanine (N-MUT, R88A/R89A/R92A/R93A; C-MUT, K256A/K257A/K261A/R262A; Figure 5A). Introduction of N-MUT and/or C-MUT into full-length (FL) or truncated N-protein clearly impaired KPNA2 interaction (Figure 5B). Co-precipitation of KPNB1 was also notably reduced by N-MUT mutations, while the effect of C-MUT on KPNB1 association was more variable; C-MUT clearly reduced KPNB1 co-precipitation with GFP-N_FL_, but had minimal effect with GFP-N_175-419_ and GFP-N_248-419_. Nevertheless, the data indicate that both sites in N-protein contribute to KPNA2 and KPNB1 association.

### SARS-COV-2 N-protein has functionally distinct associations with KPNB1 and KPNA2

If KPNA/KPNB mediate nuclear import of N-protein *via* the NTD and CTD NLS-like sequences, we would expect the mutations to also reduce the nuclear localisation of N-proteins. However, the mutations largely did not affect the steady state Fn/c of N-proteins/fragments, and even caused a modest increase in nuclear localisation in some cases (Figure 6A-C). This was most apparent with the CTD-C-tail fragment (GFP-N_248-419_), whereby introduction of C-MUTs clearly enhanced nuclear localisation (Figure 6A) and increased Fn/c (Figure 6B). Thus, reduced KPNA2/KPNB1 binding does not appear to impair N-protein nuclear import. Interestingly, GFP-N_1-174_ clearly accumulated in nucleoli (Figure S4A), and this was impaired by N-MUT (Figure 6A), suggesting that residues R88/R89/R92/R93 can drive nucleolar interactions. The lack of nucleolar accumulation by GFP-N_FL_ therefore suggests that nucleolar localisation is inhibited or regulated in the full-length N-protein.

**Figure 6.**
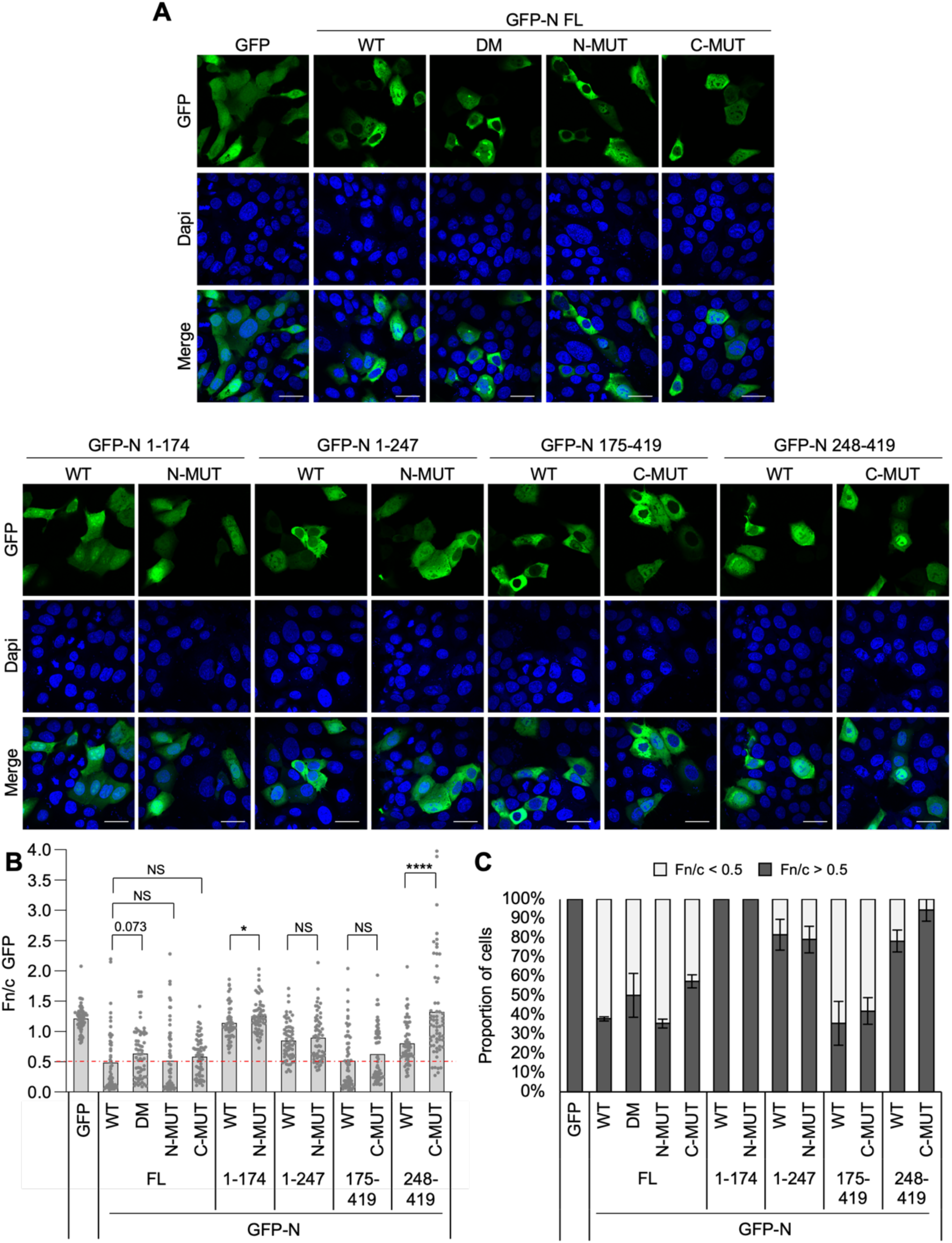
Mutation to NLS-like sequences in the CTD and NTD of SARS-COV-2 N-protein does not impair nuclear localisation. (A-C) HCT-8 cells were transfected to express the indicated proteins for 16 h before fixation and CLSM (A) to determine the Fn/c for GFP (B; mean, n = 62 cells for each condition from a representative assay) and the percentage of cells with Fn/c < or > 0.5 (C; mean ± SD, n = 2 independent assays). Dapi (blue) was used to localise nuclei. Statistical analysis used Student’s *t*-test. *, p < 0.05; ****, p < 0.0001; NS, not significant. Scale bars, 30 μm. Results are representative of two independent assays.

To confirm that the N-MUT and C-MUT mutations do not impair N-protein nuclear import, we next examined the nuclear import kinetics of the double mutant (DM) using FRAP (Figure 7A-B). Similar to WT, DM had clear ‘diffuse’ (Fn/c > 0.5) and ‘cytoplasmic’ (Fn/c < 0.5) sub-populations (Figure 7A/7C), with diffuse DM recovering markedly faster from nuclear bleaching than cytoplasmic DM (Figure 7D). Importantly, diffuse DM recovered at a similar rate (Figure 7D) and to a similar level (Figure 7E) to the analogous WT population, again indicating that the mutations do not impair nuclear import of N-protein, despite reduced KPNA2/KPNB1 binding. Interestingly, cytoplasmic DM had a significantly higher initial rate (Figure 7D) and maximal recovery (Figure 7E) than cytoplasmic WT; together with its marginally higher steady state Fn/c (Figure 6), this suggests that disabling the NLS-like sequences in the NTD and CTD may actually promote N-protein nuclear import.

**Figure 7.**
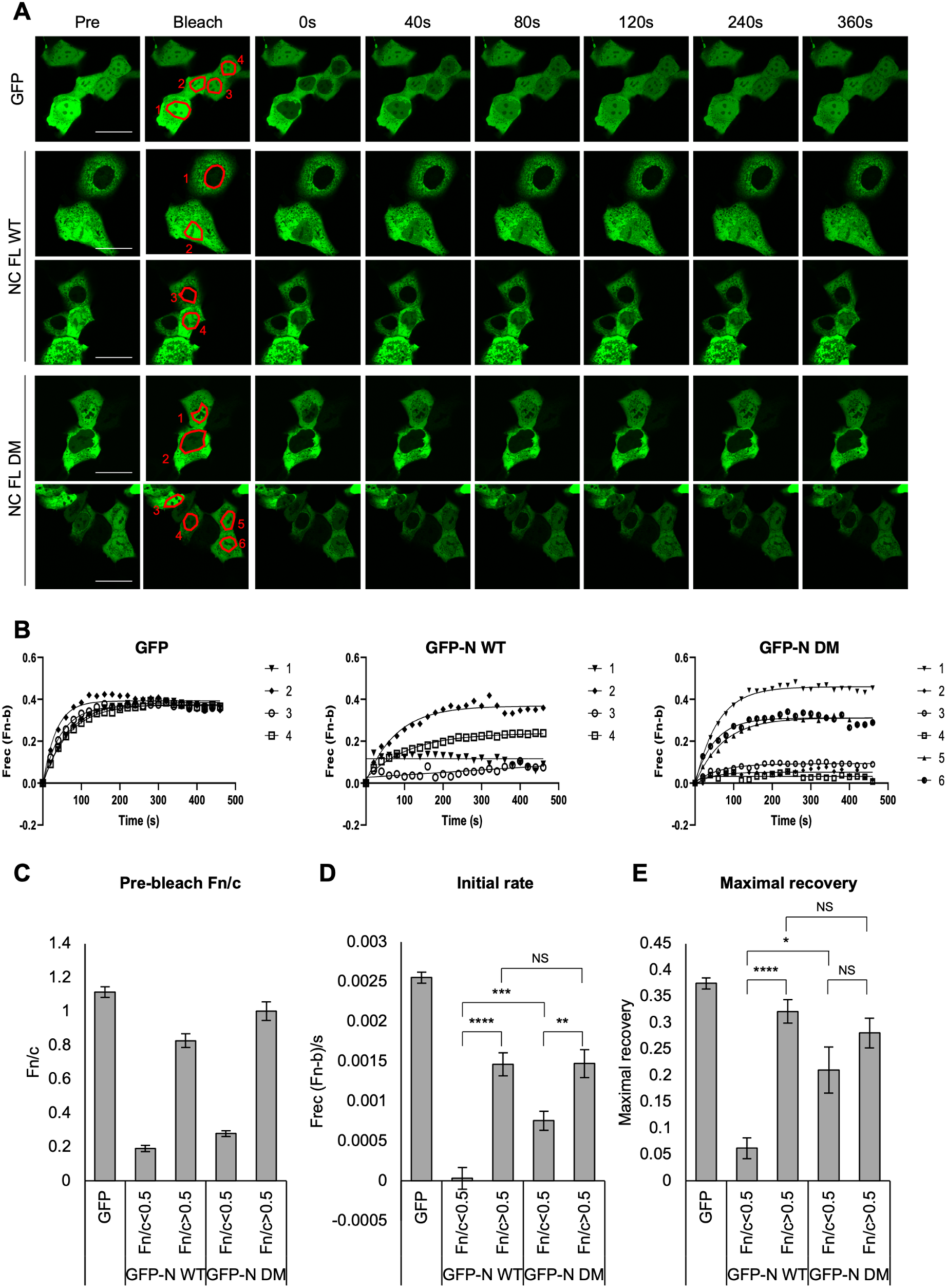
N-MUT and C-MUT do not impair nuclear import of SARS-COV-2 N-protein. (A) HCT-8 cells transfected to express the indicated proteins were imaged by CLSM prior (Pre) to photobleaching in the indicated nucleus (Bleach) and then monitored every 20s for 460s. Representative images are shown. (B) Digitised images, such as those in A, were analysed to determine the fractional change of nuclear fluorescence (Frec(Fn-b)). (C) The pre-bleach Fn/c value was determined and GFP-N samples were grouped based on Fn/c < or > 0.5. (D, E) Curves such as those in B were used to determine the initial rate of import (initial 100s post-bleaching, D) and maximal recovery of nuclear fluorescence (E). Bar charts represent the mean ± SEM (n ≥ 29 cells for each condition). Statistical analysis used Student’s *t*-test. *, p < 0.05; **, p < 0.01; ***, p < 0.001; ****, p < 0.0001; NS, not significant. Scale bars, 30 μm.

We next examined the role of the NLS-like sequences in N-protein re-localisation of KPNA2 and KPNB1. The diffuse population of WT reduced nuclear localisation of KPNA2, as above, while diffuse DM had no effect (Figure 8A-B). Importantly, this effect was not limited to KPNA2, a member of the Rch1 KPNA sub-family [30]; diffuse WT, but not DM, was also able to significantly reduce the nuclear localisation of KPNA1 (NPI-1 sub-family), and KPNA3/KPNA4 (Qip1 sub-family), albeit to differing extents (Figure S5). Thus, KPNA family re-localisation by N-protein is a) mediated by binding to NLS-like sequences in N-protein, and b) independent of N-protein nuclear import. In contrast, the mutations did not impair the capacity of diffuse N-protein to drive an increase in nuclear localisation of KPNB1 (Figure 8A-B). Thus, the effects of N-protein on KPNA and KPNB1 localisation are independent of each other, indicative of distinct functions/mechanisms. This also suggests that KPNB1 mediates nuclear import of N-protein *via* a distinct sequence/s. This could be *via* the KPNA-independent pathway, which involves KPNB1 binding directly to cargo without the KPNA adaptor [3], as treatment of cells with ivermectin, a known inhibitor of the classical KPNA-KPNB1 pathway [31], reduced nuclear localisation of GFP-T-ag_NLS_ but not GFP-N (Figure S6). Notably, the diffuse population of GFP-N_175-419_ (LKR-CTD-C-tail), but not GFP-N_1-174_ (N-tail-NTD), promoted nuclear re-localisation of KPNB1 (Figure S7), suggesting that a KPNB1-dependent NLS may be located within this C-terminal region. Consistent with this, KPNB1 was co-precipitated by GFP-N_175-419_ even in the presence of C-MUTs, albeit to a low level (Figure 5B). Finally, to confirm the role of KPNB1 in N-protein nuclear import, we depleted KPNB1 levels by siRNA knockdown (Figure 9A-B), which reduced nuclear localisation of WT and DM (Figure 9C-F). Thus, KPNB1 is a meditator of N-protein nuclear import, and this is independent of KPNA binding/mislocalisation *via* NLS-like sequences in the NTD and CTD. While knockdown of KPNA2 had no apparent effect on N-protein localisation (Figure 9C-F), consistent with KPNA-independent import of N-protein, we cannot rule out a role for KPNA2, given that knockdown was only partial (Figure 9A-B) and trafficking by KPNA family members is often highly redundant [3].

**Figure 8.**
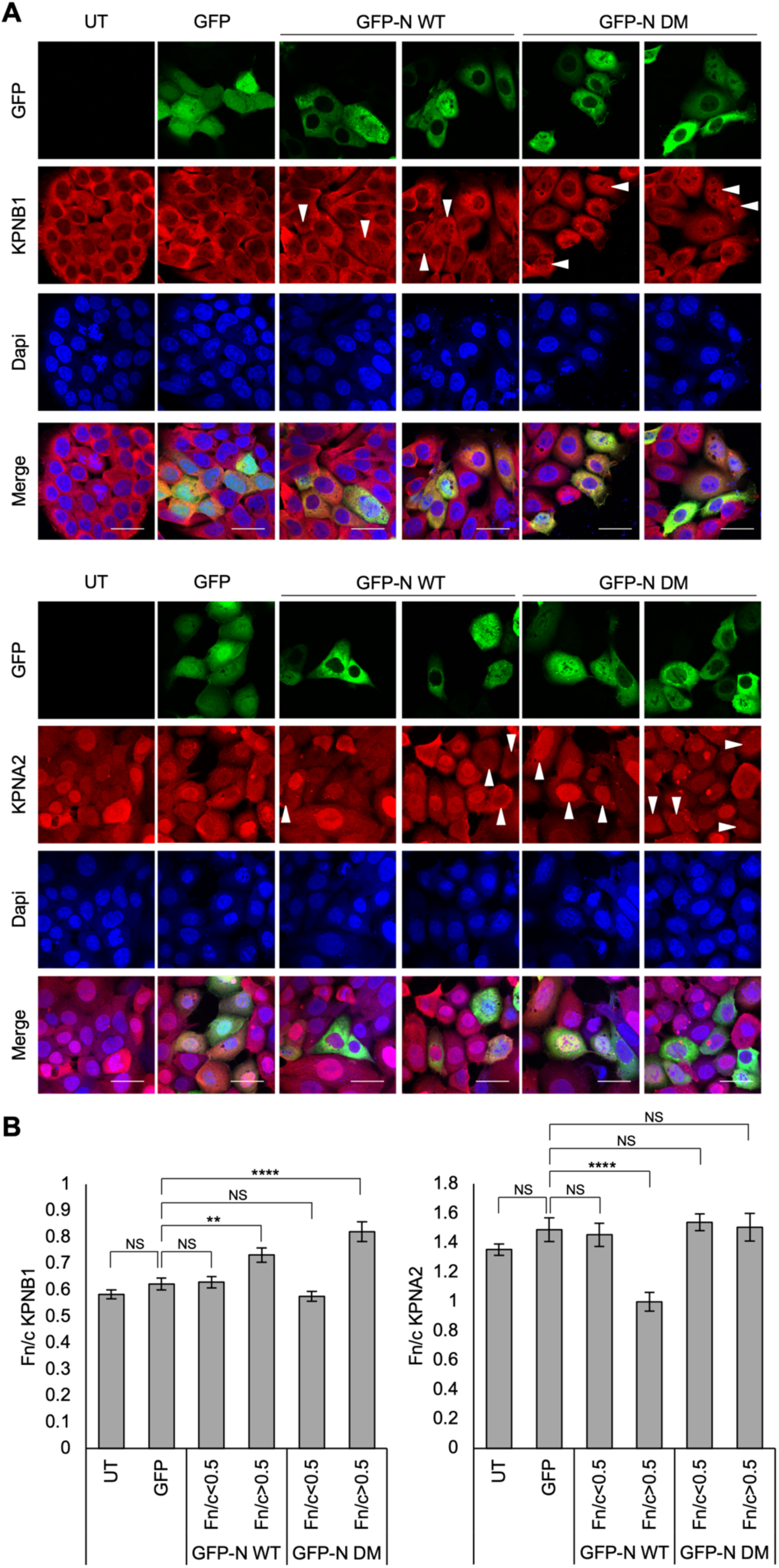
N-MUT and C-MUT impair re-localisation of KPNA2, but not KPNB1, by SARS-COV-2 N-protein. (A,B) HCT-8 cells were transfected to express the indicated proteins, or left untransfected (UT), for 16 h before fixation, immunofluorescent staining (red) for KPNB1 (upper panel) or KPNA2 (lower panel) and CLSM (A) to determine the Fn/c for KPNB1 and KPNA2 (B; mean ± SEM, n ≥ 58 cells for each condition). Only cells with clear expression of the GFP or GFP-N were analysed. GFP-N data were grouped based on GFP-N Fn/c values (Fn/c < or > 0.5). Dapi (blue) was used to localise nuclei. Arrows indicate cells with Fn/c for GFP-N > 0.5. Statistical analysis used Student’s *t*-test. **, p < 0.01; ****, p < 0.0001; NS, not significant. Scale bars, 30 μm. Results are representative of two independent assays.

**Figure 9.**
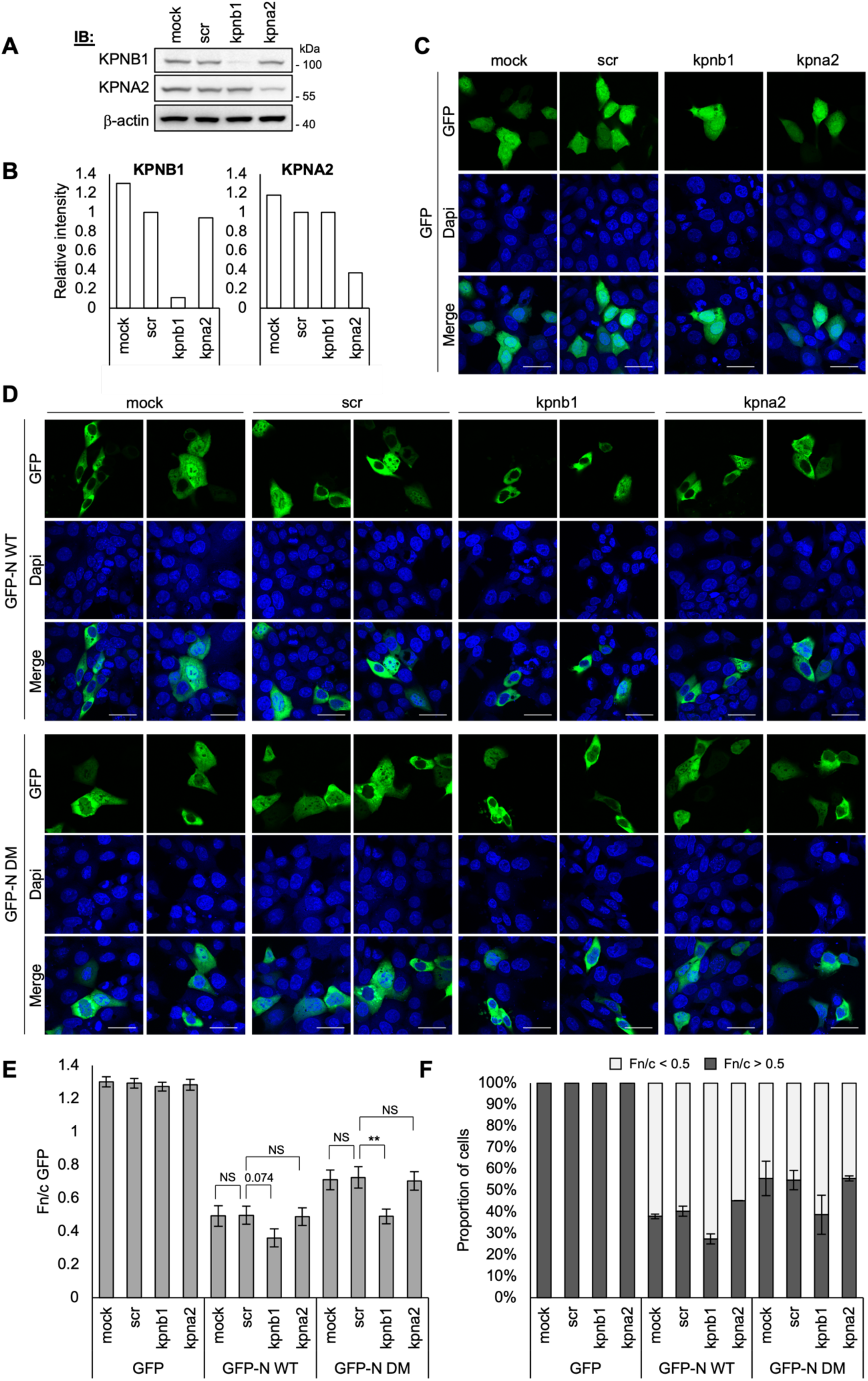
siRNA knockdown of KPNB1 reduces nuclear localisation of SARS-COV-2 N-protein. (A) HCT-8 cells were transfected with 50 nM scrambled siRNA (scr) or siRNA specific for kpna1 or kpnb1, or left untransfected (mock), for 48h before re-seeding (8h). Cells were then transfected to express the indicated proteins for 16 h before lysis and analysis by immunoblotting (IB) using antibodies against the indicated proteins. (B) Images of membranes such as those shown in (A) were analysed using Image Lab software to calculate the intensity of bands for KPNB1 and KPNA2, relative to the intensity of the corresponding β-actin band and then normalized to scrambled siRNA sample. (C-F) HCT-8 cells transfected with siRNA and treated as described in (A) were fixed and analysed by CLSM (C,D) to determine the Fn/c for GFP (E; mean ± SEM, n = 62 cells for each condition from a representative assay) and the percentage of cells with Fn/c < or > 0.5 (F; mean ± SD, n = 2 independent assays). Dapi (blue) was used to localise nuclei. Statistical analysis used Student’s *t*-test. **, p < 0.01; NS, not significant. Scale bars, 30 μm. Results are representative of two independent assays.

## Discussion

### N-protein undergoes nuclear import via KPNB1

Here we have identified distinct populations of cells expressing SARS-COV-2 N-protein, distinguishable by the steady state localisation of N-protein and the capacity for N-protein to undergo nuclear import. In most cells, N-protein is highly cytoplasmic (Figure 1) and largely incapable of nuclear entry (Figure 2), consistent with the majority of studies that describe cytoplasmic localisation of N-protein (e.g [6–10]). However, we find that in a proportion of cells, the size of which changes over time, N-protein localises more diffusely between the cytoplasm and nucleus (Figure 1), driven by nuclear import *via* a KPNB1-dependent pathway. This would explain the presence of N-protein in the nucleus reported in several studies [15–18]. Evidence supporting a role for KPNB1 in N-protein nuclear import includes 1) N-protein co-precipitates KPNB1 (Figure 3A and Figure S2A), 2) nuclear localisation of N-protein is reduced when KPNB1 is knocked down (Figure 9), and 3) N-protein effects an increase in the nuclear localisation of KPNB1 (Figure 3B-C), consistent with active N-protein-KPNB1 complexes in the nucleus. Mutation to the NLS-like sequences at residues 88-95 and 256-262, while reducing KPNB1 binding (Figure 5B), did not impair N-protein nuclear trafficking (Figure 7 and Figure 9), indicating that the ‘real’ NLS is distinct from these sites. It is worth noting that trafficking complexes are transient and so often have lower binding affinity than inhibitory complexes [32], making them difficult to detect by immunoprecipitation (see T-ag_NLS_ in Figure 3A and Figure S2A). Thus, it is feasible that the relatively small amount of KPNB1 co-precipitated by the mutants (Figure 5B) is sufficient for import. Notably, the fragment GFP-N_175-419_ is > 55 kDa, which is larger than the NPC diffusion limit, and yet, like WT, has distinct cytoplasmic and diffuse populations, with the diffuse population capable of re-localising KPNB1 to the nucleus (Figure S7). This region therefore warrants further investigation for a KPNB1-dependent NLS.

N-protein’s capacity to be imported into the nucleus suggests intranuclear functions, distinct from its well-established cytoplasmic roles in genome replication and virus assembly. This may relate to the cell cycle, as is the case with nucleolar IBV N-protein [5,24]. Interestingly, SARS-COV-2 N-protein was recently shown to associate and co-localise with nucleolin in nucleoli, and re-localise nucleolin to the cytoplasm [15]. Nucleolin appears to be pro-viral for SARS-COV-2, contributing to virus-induced apoptosis [15]. While full-length SARS-COV-2 N-protein did not accumulate in nucleoli in our assays, consistent with several other studies [6–10], the fragment GFP-N_1-174_ did, and this was dependent on the NLS-like sequence at residues 88-95 (Figure 6 and Figure S4). Notably, this is distinct from the region in SARS-COV-1 N-protein that is reported to mediate nucleolar localisation (residues 226-289) [13,14]. The lack of nucleolar localisation of full-length N-protein here, and in other studies, suggests the presence of other binding sites and/or sequences preventing nucleolar localisation, which are absent or disrupted in GFP-N_1-174_. It is also possible that nucleolar localisation of full-length N-protein requires other SARS-COV-2 proteins that are absent in our system. It will therefore be interesting to examine what conditions are required to ‘activate’ nucleolar accumulation of SARS-COV-2 N-protein.

Viral proteins localise to the nucleus for many reasons/functions. SARS-COV-2 nsp16 was recently found to localise to the nucleus to regulate host mRNA splicing, contributing to suppression of antiviral interferon responses [33]. Interestingly, mass spectrometry analysis of the SARS-COV-2 N-protein interactome reported significant enrichment of the gene ontology processes ‘snoRNA binding’ and ‘spliceosomal complex’ among N-protein interactors [34]. It is therefore interesting to speculate that nuclear N-protein may also contribute to modulation of host RNA metabolism pathways. Moreover, preliminary analysis of a panel of SARS-COV-2 proteins found that several proteins localise to the nucleus and interact with KPNA2 and/or KPNB1, including ORF9b, nsp5 and nsp16 (Figure S2). Determining the nuclear roles of N-protein and other SARS-COV-2 proteins will be the focus of future work.

Our FRAP data suggest that the predominant ‘cytoplasmic’ population of N-protein is largely not imported into the nucleus, with minimal fluorescence recovery in the nucleus after photobleaching (Figure 2D-E). While this may seem unsurprising or expected, many ‘cytoplasmic’ viral proteins actually shuttle dynamically in and out of the nucleus, with their steady-state cytoplasmic localisation a result of the nuclear export rate exceeding that of nuclear import (e.g. Ebola virus VP24 [21]). Rather, our data indicate that only a portion of SARS-COV-2 N-protein is able to traffic to the nucleus, and this portion changes over time (Figure 1). Thus, nuclear import of N-protein appears tightly regulated and context-dependent. Given the critical roles for N-protein in encapsidating viral RNA, genome replication and virus assembly, all of which occur exclusively in the cytoplasm, it appears logical that only a proportion of N-protein is trafficked to the nucleus, and thus away from these important functions. Moreover, these functions require N-protein to be tightly wound around RNA in high-order oligomers; it is therefore possible that this population of N-protein is inaccessible to the nuclear transport machinery. Indeed, the presence of a highly cytoplasmic population required both the LKR and CTD (Figure S4), which have been implicated in N-protein oligomerisation and dimerisation, respectively [35,36]. How N-protein nuclear trafficking is regulated thus requires further investigation.

### N-protein mislocalises KPNAs via NLS-like sequences

While N-protein appears to use KPNB1 for nuclear import, we identified a functionally distinct association with KPNA. If KPNA also contributed to N-protein import, we would likely expect N-protein expression to also promote nuclear localisation of KPNA, as observed with KPNB1 (Figure 3B-C), and as seen in cells expressing GFP-T-ag_NLS_ (Figure S3), which contains a characterised KPNA/KPNB1-dependent NLS. Instead, the diffuse population of N-protein reduced the nuclear localisation of KPNA1, KPNA2, KPNA3 and KPNA4 (Figure 3B-C and Figure S5), suggesting possible antagonism (Figure S3). Moreover, ivermectin, which inhibits the activity of the KPNA-KPNB1 heterodimer and reduces nuclear localisation of GFP-T-ag_NLS_ ([31] and Figure S6), had no effect on N-protein localisation, providing further evidence that N-protein may not require KPNA for import.

We identified two NLS-like sequences (residues 88-95 and 256-262) that mediate binding to KPNA2; mutating these sites disabled KPNA2 binding (Figure 5B) and mislocalisation (Figure 8), but, importantly, did not impair nuclear import (Figure 7) or re-localisation of KPNB1 (Figure 8). Thus, KPNA2 binding/mislocalisation are independent of KPNB1-mediated import. In fact, the mutations induced a significant increase in N-protein nuclear entry (Figure 7D-E), suggesting that disrupting N-protein-KPNA2 interaction may make N-protein more available for import by KPNB1.

A number of viral proteins interact with KPNAs to inhibit the nuclear import of host cargo, and thereby supress antiviral immune responses. Ebola VP24, for example, binds competitively to KPNAs to block the nuclear import of STAT1 [32] and STAT3 [23], key mediators of interferon and interleukin-6 signalling, respectively. ORF6 from SARS-COV-1 and SARS-COV-2 also bind KPNAs and block STAT1 trafficking [19,20]. Importantly, SARS-COV-2 ORF6 appears to dysregulate nuclear trafficking more broadly by binding NPC components Rae1 and Nup98 [27,28], while other SARS-COV-2 proteins nsp1, nsp9 and nsp15 also disrupt nucleocytoplasmic transport by targeting NXF1-NXT1, Nup62 and KPNA1, respectively [37–39]. Thus, N-protein-KPNA interaction and mislocalisation may contribute to a wider dysregulation of host trafficking by SARS-COV-2. This may explain why the KPNA2-N-protein interaction appears more robust than KPNA2-T-ag_NLS_ (Figure 3A and Figure S2A); trafficking complexes are mostly transient while inhibitory complexes can have very high affinities [32]. However, it is important to note that the effect of N-protein on KPNA localisation is only partial; there is still a substantial level of KPNA in the nucleus of cells expressing diffuse N-protein, and the cytoplasmic population of N-protein does not cause a reduction in nuclear KPNA (Figure 3B-C and Figure S5). This is consistent with other SARS-COV-2 proteins being the key mediators of antagonising nuclear trafficking (above), but suggest that N-protein might provide additional fine-tuning.

We therefore propose a model in which a proportion of N-protein is capable of accessory functions, including trafficking to the nucleus and mislocalisation of KPNAs. This may be particularly important early in infection when viral proteins are translated from incoming viral RNA before genome replication has begun, such that there is likely to be substantial N-protein that is not yet encapsidating nascent viral RNA and so free to participate in a unique set of interactions and functions, including with KPNB1 and KPNA2. Given that N-protein is the most abundantly expressed viral protein in infected cells [40], even this small proportion of N-protein is likely to have notable impact. Importantly, nuclear transport inhibitors, such as ivermectin and leptomycin B, can inhibit replication by diverse viruses [2], including SARS-COV-2 [41], indicating that the interactions identified here may have value for anti-SARS-COV-2 drug development.

## Materials and Methods

### Plasmid constructs

Plasmids encoding full-length or truncated proteins from SARS-COV-2 (clinical isolate Australia/VIC01/2020) tagged at the N-terminus with eGFP were generated by GenScript in pcDNA 3.1-N-eGFP. Mutagenesis to NLS-like sequences in N-protein was performed by GenScript. Nsp1 mutations K164A/H165A were previously shown to disable nsp1-mediated host gene shut-off [42]. pEGFP-C1 plasmid encoding the NLS from the SV40 T-ag (resides 111-135) has been described elsewhere [25,26]. Plasmid to express FLAG-tagged KPNA2 was a kind gift from C. Basler (Icahn School of Medicine at Mount Sinai).

### Cells, transfections and drug treatments

HCT-8 cells were maintained in RPMI-1640 supplemented with 10% horse serum, 2 mM glutamine and 1 mM sodium pyruvate, at 5% CO_2_, 37°C. DNA and siRNA transfections used Lipofectamine 3000 (Thermo Fisher Scientific) and DharmaFECT (Horizon), respectively, according to the manufacturers’ instructions. siRNA experiments used ON-TARGETplus siRNA SMARTpool (Dharmacon) targeting human KPNB1 (L-017523-00-0005) or KPNA2 (L-004702-00-0005). For negative control we used ON-TARGETplus Non-targeting Pool (Dharmacon, D-001810-10-50). siRNA was transfected at a final concentration of 50 nM for 48 h; cells were then re-seeded (8 h) and transfected with plasmid DNA for 16 h before downstream analysis. To inhibit the KPNA/KPNB1 nuclear import pathway, cells were treated with 25 µM ivermectin or DMSO control for 1.5 h.

### CLSM for steady state localisation

Cells pre-seeded onto coverslips and transfected/treated as specified were fixed with 4% PFA for 20 min at 37°C, permeabilised with PBS/0.02% Triton-X-100 for 20 min, blocked with PBS/1% BSA for 1 h and immunostained at 4°C overnight with primary antibodies diluted in blocking buffer. Antibodies used were: anti-KPNB1 (Abcam, ab2811), anti-KPNA1 (Abcam, ab154399), anti-KPNA2 (BD Biosciences, 610486), anti-KPNA3 (Abcam, ab6038) and anti-KPNA4 (Abcam, ab6039). Coverslips were then washed five times with PBS before staining with Alexa Fluor 568 secondary antibodies (Thermo Fisher Scientific) at RT for 1 h, and mounting using Prolong-gold antifade mounting reagent with DAPI (Thermo Fisher Scientific). Imaging used a Leica SP5 or Nikon C1 Inverted CLSM with 63 X objective. Digitised images were processed using Fiji software (NIH). To quantify nucleocytoplasmic localisation, the ratio of nuclear to cytoplasmic fluorescence (Fn/c) was calculated for individual cells using the formula Fn/c = (Fn-Fb)/(Fc-Fb), where Fn is the nuclear fluorescence, Fc is the cytoplasmic fluorescence and Fb is the background autofluorescence, as previously [21–23]. The mean Fn/c was then calculated for n > 35 cells.

### FRAP to analyse nuclear import

Transfected cells were imaged live in phenol-free media using an Olympus Fluoview 1000 CLSM with a 100X oil immersion objective at 37°C. As previously [43,44], the nucleus was photobleached with high laser power (12 scans, 20 s/pixel) and the fluorescence recovery in the nucleus was monitored with low laser power (12.5 s/pixel) at 20 s intervals for 460 s. Data are presented as fractional recovery (Frec), which reflects the nuclear fluorescence (Fn) minus background fluorescence (Fb), relative to the pre-bleached values (Fn_pre), calculated at each time point according to the formula: Frec(Fn − b) = (Fn − Fb)/Fn_pre. The initial rate of recovery (Frec (Fn − b)/s) was calculated by linear regression of data points from the first 100 s, where the rate of fluorescence recovery was approximately linear. The maximal level of fluorescence recovery (maximal recovery) was determined by plotting individual values for Frec(Fn − b) (y) against time (t) to a nonlinear equation in Prism 7 [43,44].

### Immunoprecipitations and western blotting

Transfected cells were lysed for immunoprecipitation using GFP-Trap beads (Chromotek), according to the manufacturer’s instructions. Lysis and wash buffers were supplemented with cOmplete Protease Inhibitor Cocktail (Roche). Lysates and immunoprecipitates were analysed by SDS-PAGE and immunoblotting using antibodies against KPNA2 (BD Biosciences, 610486), KPNB1 (BD Biosciences, 610560), GFP (Roche, 11814460001 and Abcam, ab290), β-actin (Cell Signaling Technology, 3700), and HRP-conjugated secondary antibodies (Merck). Visualization of bands used Western Lightning chemiluminescence reagents (PerkinElmer) and ChemicDoc Imaging Systems (Bio-Rad). Image Lab software (Bio-Rad) was used to calculate the relative intensity of bands.

### Statistical Analysis

Unpaired two-tailed Student’s t-test was performed using Microsoft Excel and Prism software (version 7, GraphPad).

## Data Availability

Data available upon request to Angela R. Harrison: or Kylie M. Wagstaff

## Acknowledgements

We thank the Monash Micro Imaging Facility for assistance with CLSM.

---

**Supplemental Figure S1.**
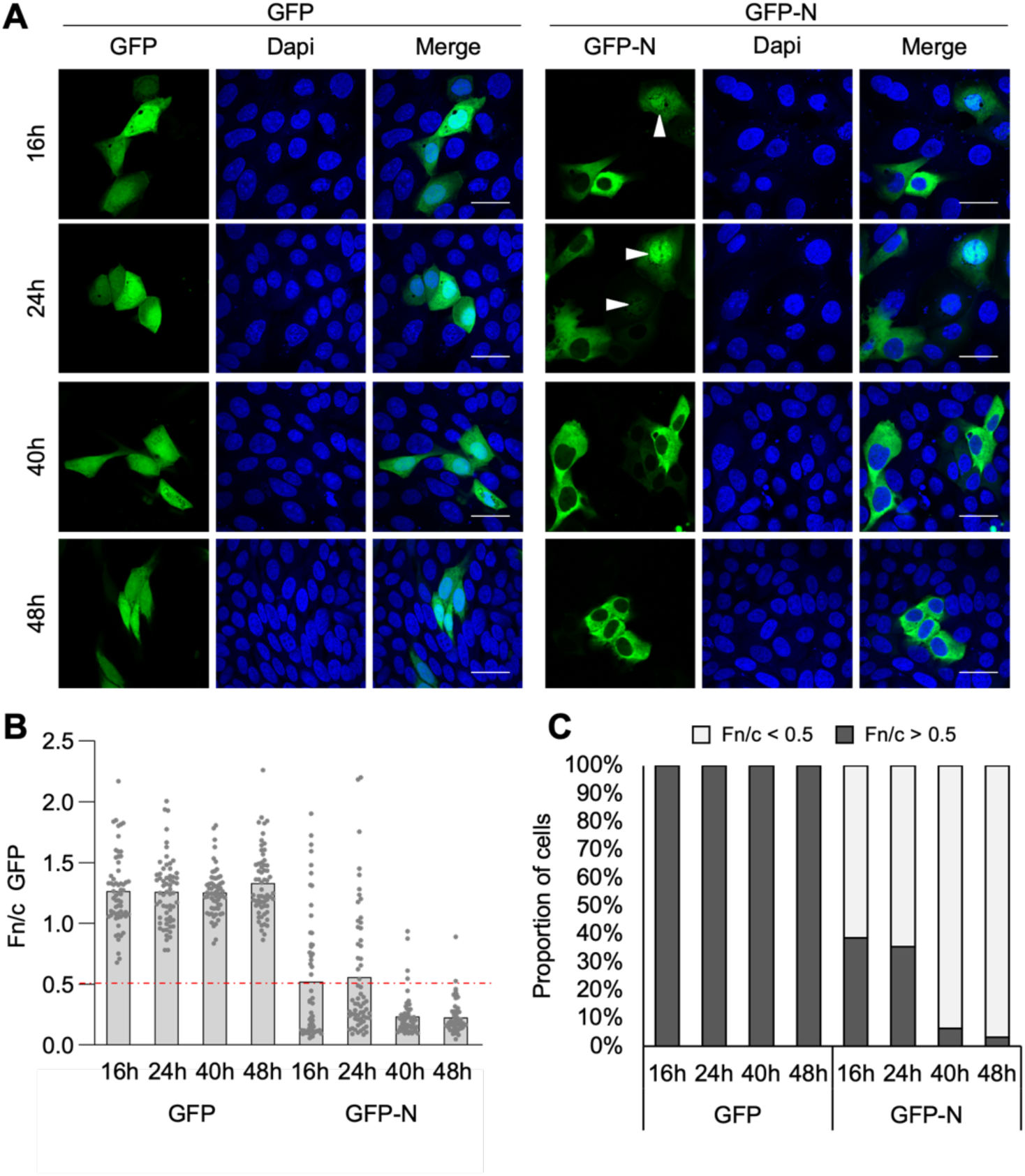
Distinct populations of SARS-COV-2 N-protein are present in synchronised cells. (A) HCT-8 cells were starved for 24 h in serum-free media to synchronise cells, before replacement with complete media and transfection to express GFP or GFP-N for the indicated time. Cells were then fixed for CLSM analysis. Representative images are shown. (B,C) Images such as those in (A) were analysed to calculate the Fn/c for GFP (B; mean, n = 62 cells for each condition) and the percentage of cells with Fn/c < or > 0.5 (C). Dapi (blue) was used to localise nuclei. Arrows indicate cells with Fn/c for GFP-N > 0.5. Scale bars, 30 μm.

**Supplemental Figure S2.**
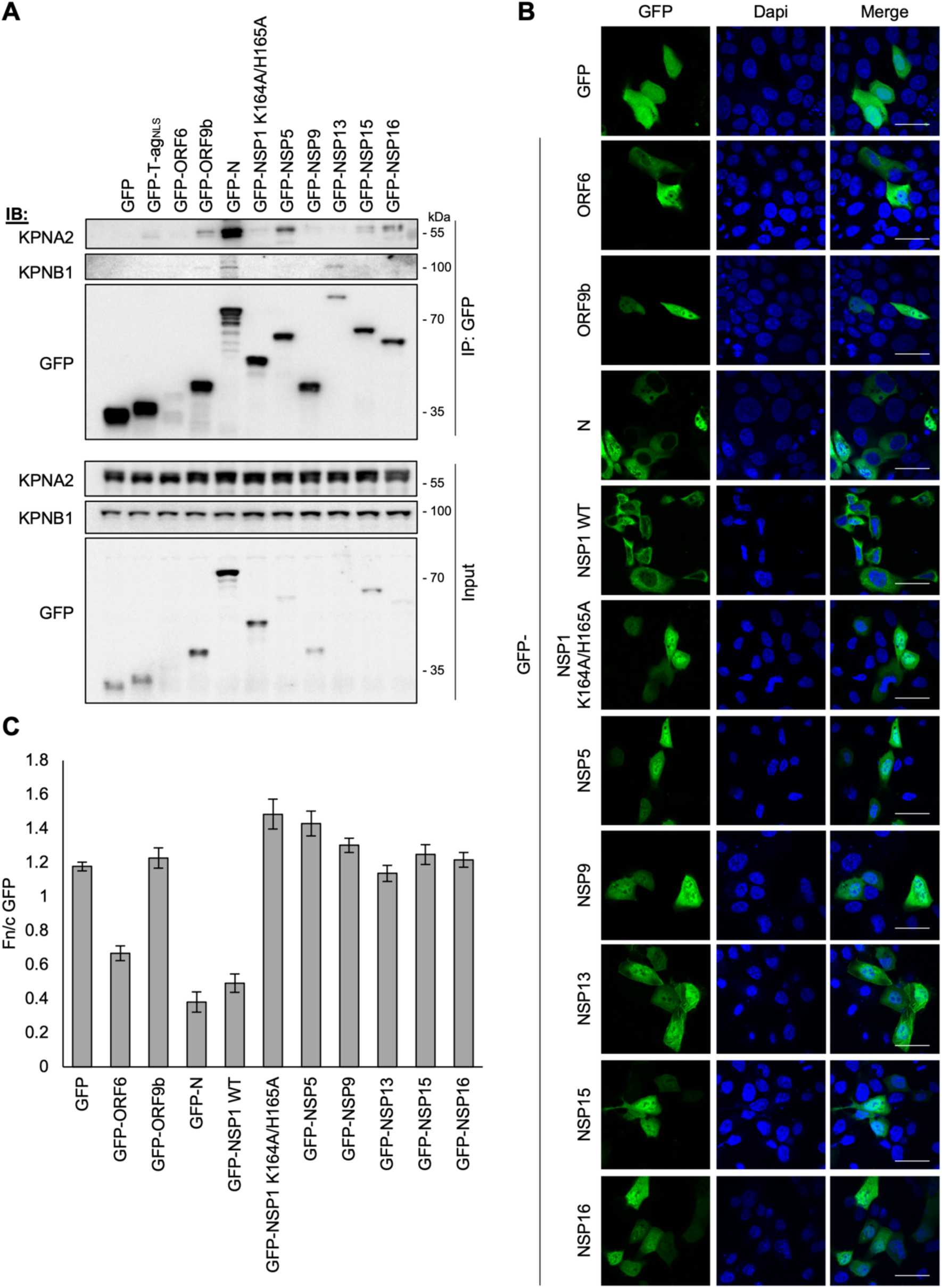
Several SARS-COV-2 proteins co-precipitate KPNB1 and/or KPNA2 and are present in the nucleus. (A) HCT-8 cells were co-transfected to express GFP or the indicated GFP-tagged protein and FLAG-tagged KPNA2 for 24 h before lysis and immunoprecipitation for GFP. Lysates (Input) and immunoprecipitates (IP) were analysed by immunoblotting (IB) using antibodies against the indicated proteins. (B-C) HCT-8 cells were transfected to express the indicated proteins for 16 h before fixation and CLSM (B) to determine the Fn/c for GFP (C; mean ± SEM, n ≥ 39 cells for each condition). Dapi (blue) was used to localise nuclei. Scale bars, 30 μm. Results are representative of two independent assays.

**Supplemental Figure S3.**
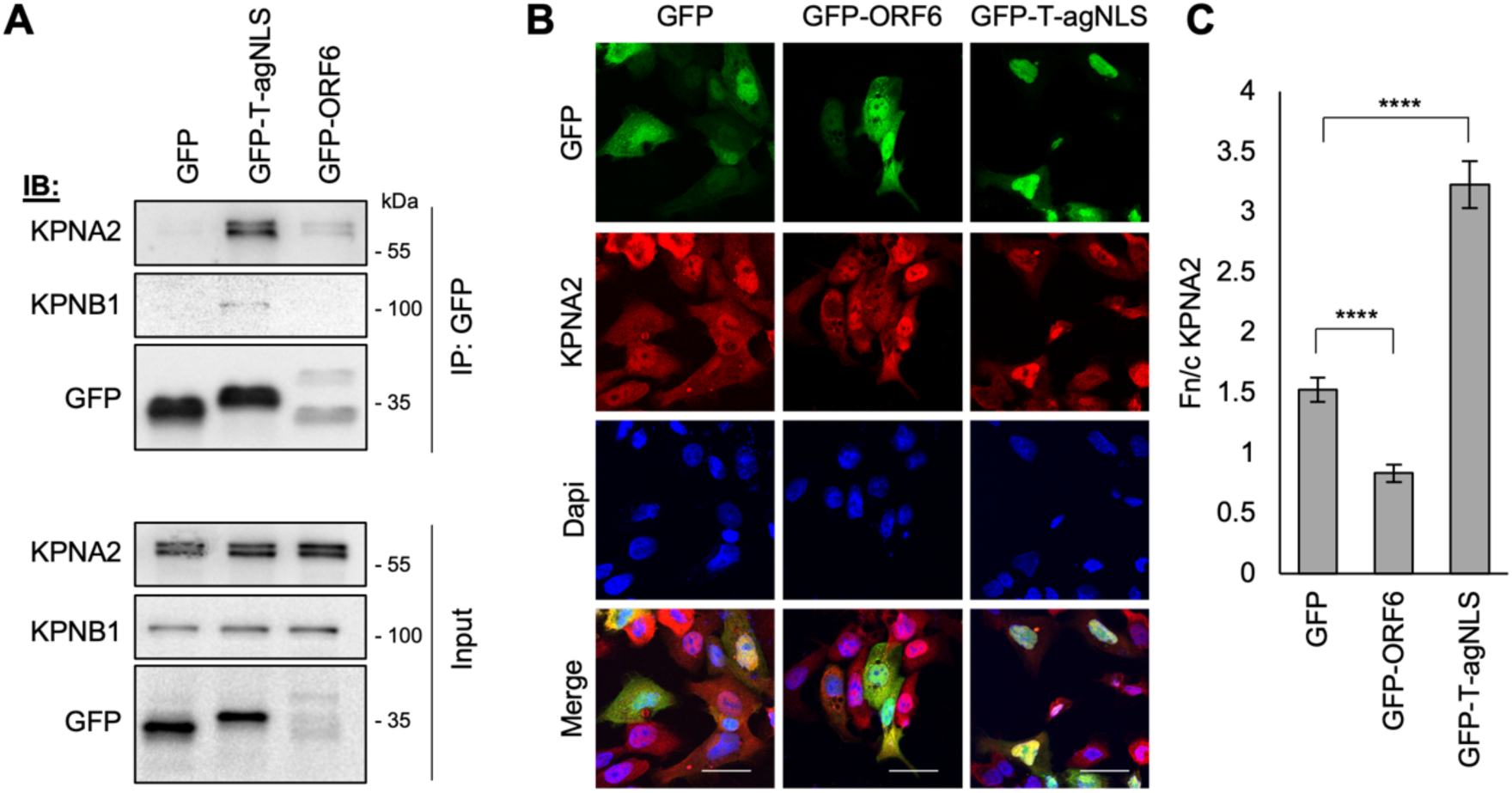
T-ag_NLS_ and SARS-COV-2-ORF6 target KPNA2 for distinct functions. (A) HCT-8 cells were co-transfected to express GFP, GFP-T-ag_NLS_ or GFP-ORF6 and FLAG-tagged KPNA2 for 24 h before lysis and immunoprecipitation for GFP. Lysates (Input) and immunoprecipitates (IP) were analysed by immunoblotting (IB) using antibodies against the indicated proteins. (B,C) HCT-8 cells were transfected to express GFP, GFP-T-ag_NLS_ or GFP-ORF6 for 16 h before fixation, immunofluorescent staining for KPNA2 (red) and CLSM (B) to determine the Fn/c for KPNA2 (C; mean ± SEM, n ≥ 52 cells for each condition). Only cells with clear expression of GFP or GFP-tagged protein were analysed. Dapi (blue) was used to localise nuclei. Statistical analysis used Student’s *t*-test. ****, p < 0.0001. Scale bars, 30 μm.

**Supplemental Figure S4.**
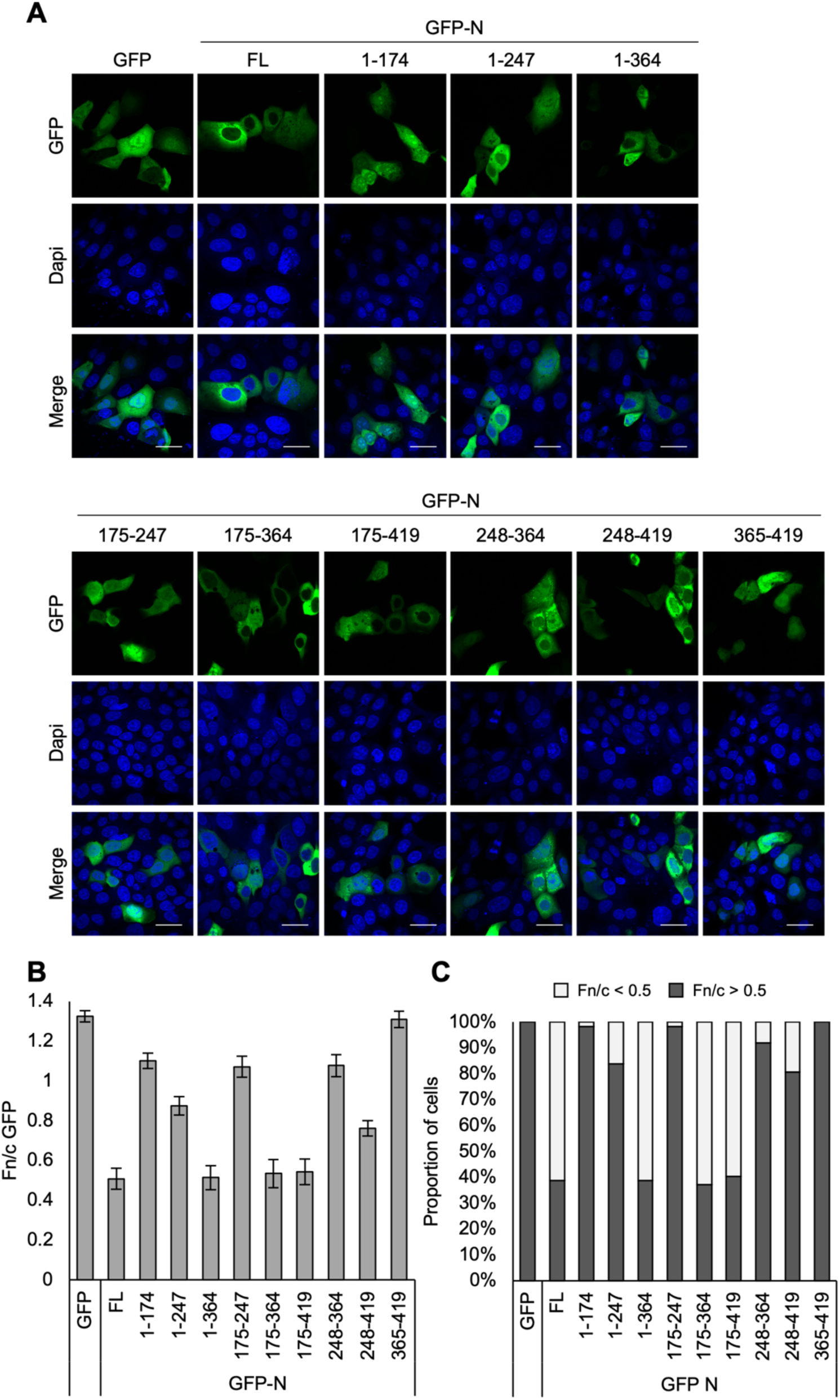
Truncation of SARS-COV-2 N-protein affects nucleocytoplasmic localisation. (A-C) HCT-8 cells were transfected to express the indicated proteins for 16 h before fixation and CLSM (A) to determine the Fn/c for GFP (B; mean ± SEM, n = 62 cells for each condition) and the percentage of cells with Fn/c < or > 0.5 (C). Dapi (blue) was used to localise nuclei. Scale bars, 30 μm.

**Supplemental Figure S5.**
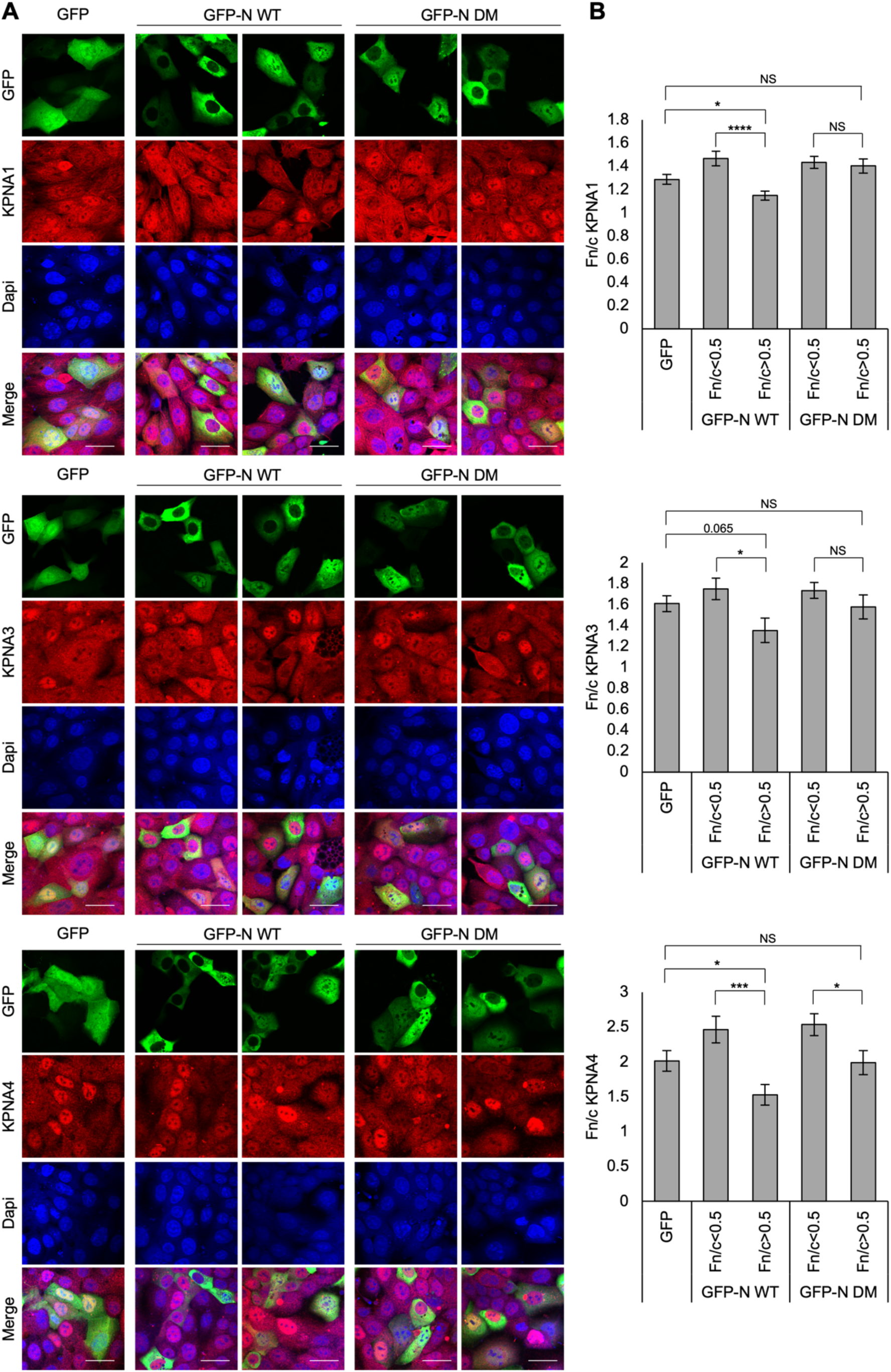
SARS-COV-2 N-protein re-localises multiple KPNAs, dependent on NLS-like sequences. (A,B) HCT-8 cells were transfected to express the indicated proteins for 16 h before fixation, immunofluorescent staining (red) for KPNA1 (upper panel), KPNA3 (middle panel) or KPNA4 (lower panel) and CLSM (A) to determine the Fn/c for KPNA1, KPNA3 and KPNA4 (B; mean ± SEM, n ≥ 60 cells for each condition). Only cells with clear expression of GFP or GFP-N were analysed. GFP-N data were grouped based on GFP-N Fn/c values (Fn/c < or > 0.5). Dapi (blue) was used to localise nuclei. Statistical analysis used Student’s *t*-test. *, p < 0.05; ***, p < 0.001; ****, p < 0.0001; NS, not significant. Scale bars, 30 μm.

**Supplemental Figure S6.**
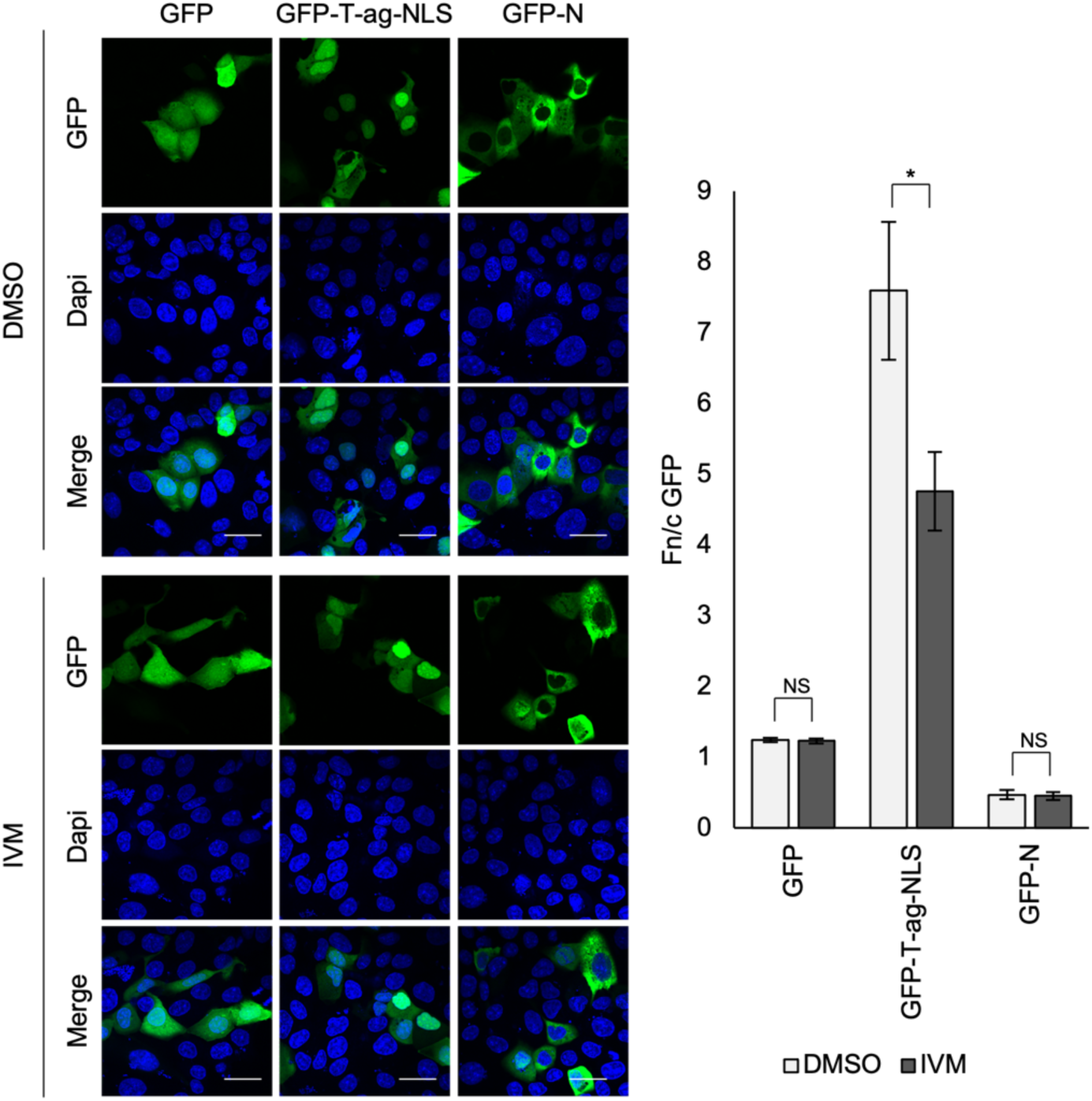
The nucleocytoplasmic localisation of SARS-COV-2 N-protein is not affected by ivermectin treatment. (A-B) HCT-8 cells transfected to express the indicated proteins were treated with 25µM ivermectin (IVM) or DMSO control for 1.5 h before fixation and CLSM (A) to determine the Fn/c for GFP (B; mean ± SEM, n = 62 cells for each condition). Dapi (blue) was used to localise nuclei. *, p < 0.05; NS, not significant. Scale bars, 30 μm. Results are representative of two independent assays.

**Supplemental Figure S7.**
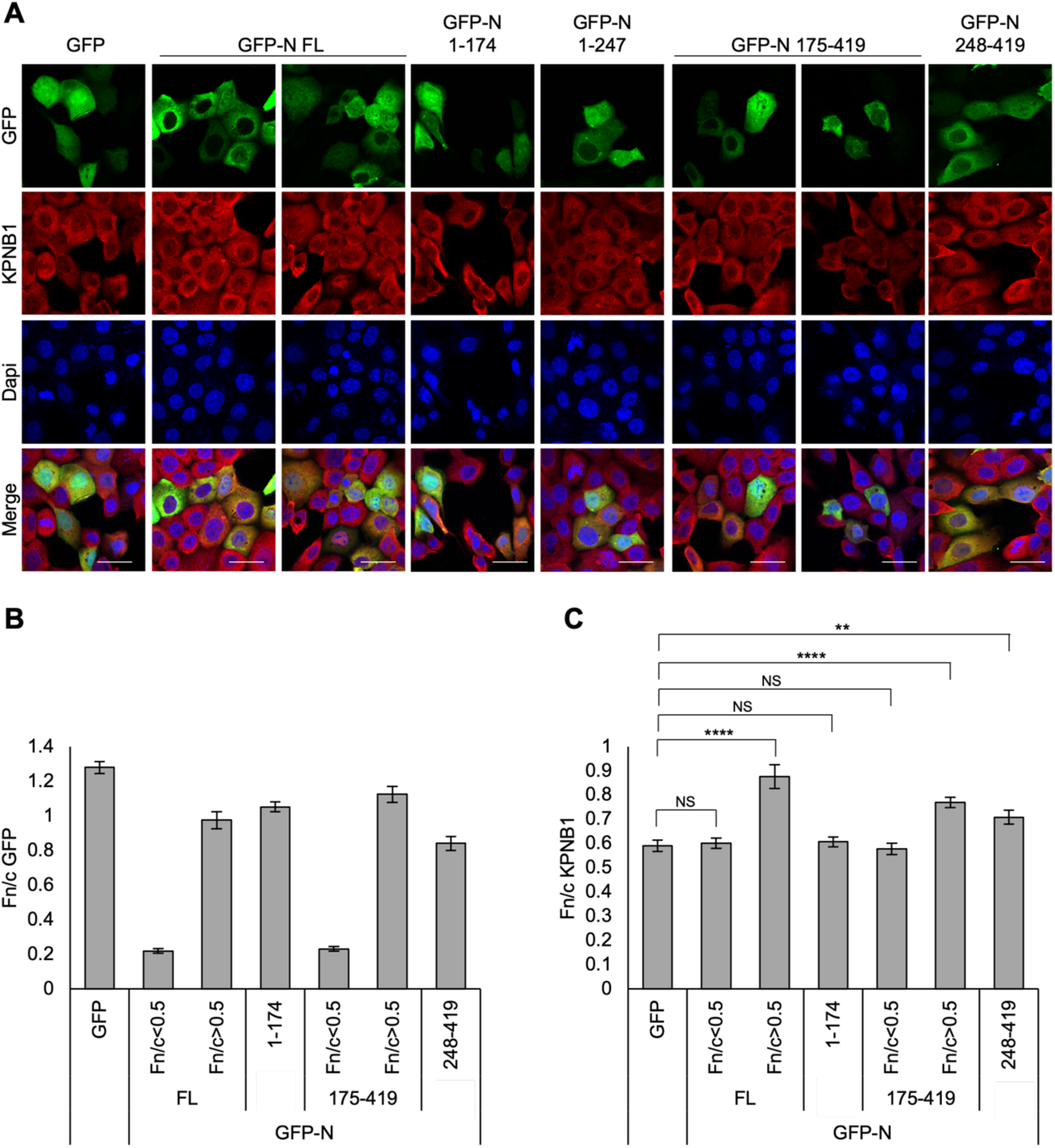
Re-localisation of KPNB1 by SARS-COV-2 N-protein requires the CTD-C-tail. (A-C) HCT-8 cells were transfected to express the indicated proteins for 16 h before fixation, immunofluorescent staining for KPNB1 (red) and CLSM (A) to determine the Fn/c for GFP (B) and KPNB1 (C; mean ± SEM, n = 62 cells for each condition). Only cells with clear expression of GFP or the GFP-tagged protein were analysed. GFP-N FL and 175-419 data were grouped based on GFP-N Fn/c values (Fn/c < or > 0.5). Dapi (blue) was used to localise nuclei. Statistical analysis used Student’s *t*-test. ****, p < 0.0001; **, p < 0.01; NS, not significant. Scale bars, 30 μm.

